# One-shotted: Psychedelic insights are uniquely intense, meaningful, and ineffable

**DOI:** 10.64898/2026.06.02.729484

**Authors:** Ambra Pogliani, Alexandra Zachary, Lena Hall, Jason M. Tangen, Jonathan Mond, Ruben E. Laukkonen

## Abstract

Insight is commonly studied in laboratory paradigms as a sudden shift in understanding accompanied by characteristic “Aha!” experiences. However, such paradigms may not capture the full phenomenological range of insight as it occurs in naturalistic contexts. This study compared the phenomenology of insight across four contexts—psychedelic experiences, everyday-life insights, Compound Remote Associates problems, and ambiguous images—in a within-subject online study of adults with prior psychedelic experience (N = 73). Participants rated each insight for intensity, surprise, confidence, pleasure, drive, meaning, ineffability, and perceived belief change. Ordinal mixed-effects models showed that psychedelic insights were rated higher than everyday insights on most dimensions, especially intensity, meaning, ineffability, and perceived belief change, whereas laboratory insights were substantially lower than both naturalistic contexts. Naturalistic–laboratory differences were largest for meaning and perceived belief change and smaller for core “Aha!” features such as confidence and pleasure. Within naturalistic contexts, meaning was the strongest predictor of perceived belief change, with additional contributions from intensity and ineffability; after these phenomenological dimensions were included, context no longer predicted perceived belief change. These findings suggest that laboratory paradigms capture core features of insight but underrepresent dimensions related to personal meaning and belief updating. They further indicate that insight-related belief change depends less on context itself than on how the insight is experienced.

## 1. INTRODUCTION

In the field of cognitive psychology, insight is commonly defined as a sudden shift in understanding, where a stimulus, situation, or problem is re-interpreted in a novel way (Kounios & Beeman, 2015). Although insight refers to this cognitive change in interpretation, it is often accompanied by a distinctive subjective experience—the “Aha!” or “Eureka” moment—characterised by feelings of sudden clarity, surprise, perceived rightness, and affective salience (Topolinski & Reber, 2010; Wiley & Danek, 2024). In the present paper, these are treated as related but distinguishable aspects of the broader phenomenon: insight refers to the cognitive event (cognitive restructuring that leads to a solution or to an idea), whereas “Aha!” phenomenology refers to the subjective feelings that accompany it, and may help signal the emergence of that idea.

In cognitive psychology, insight has been studied most extensively using laboratory paradigms designed to reliably elicit brief “Aha!” experiences under controlled conditions. A widely used example is the Compound Remote Associates (CRA) task, in which participants identify a word that links three seemingly unrelated words (Bowden & Jung-Beeman, 2003; Chein & Weisberg, 2014; Webb et al., 2018). Insight has also been studied using perceptual paradigms, such as ambiguous or bistable images, where an “Aha!” moment arises from a sudden reorganisation of perceptual interpretation (Kornmeier & Bach, 2012; Schooler & Melcher, 1995). Evidence suggests that verbal and perceptual insight share meaningful similarities: individuals who perform well on ambiguous images also tend to perform well on insight problems, and both are accompanied by the characteristic “Aha!” phenomenology (Doherty & Mair, 2012; Laukkonen & Tangen, 2018). Together, these and other laboratory paradigms have enabled researchers to characterise a recurring phenomenological profile of insight, including surprise, confidence, sense of truth, positive affect (e.g., pleasure or satisfaction), and motivational drive (Danek et al., 2014; Danek & Wiley, 2017). However, these tasks have limited ecological validity as they rely on constrained problems with a single correct solution and typically carry little personal relevance. As a result, they are well suited to capturing the core “Aha!” signal (e.g., confidence and surprise), but may not fully reflect context-sensitive qualities of insight in naturalistic settings where experiences are often personally salient and affectively significant, and may therefore be experienced as more meaningful, intense, and impactful for beliefs and behaviour (Hill & Kemp, 2018; Jennissen et al., 2018; Tulver et al., 2023).

A growing body of work has begun to examine insight beyond the laboratory, including everyday-life and clinical contexts. In everyday settings, people report insights ranging from practical problem-solving solutions to personally meaningful realisations about themselves and their lives (Hill & Kemp, 2018). In psychotherapy, insight commonly involves recognising links between past and present experience, identifying relational or emotional patterns, and understanding their impact on psychological symptoms, behaviour and psycho-social functioning (Lacewing, 2014). Importantly, meta-analytic evidence suggests that such insights are positively associated with psychotherapy outcomes (Jennissen et al., 2018). These findings indicate that insight is not limited to brief, puzzle-like solutions and may be particularly important when it reorganises higher-level interpretations that guide emotion regulation, self-concept, and behaviour.

These issues become especially salient in psychedelic contexts, that is, during or following the acute effects of psychedelic substances (e.g. psilocybin, LSD). Insights arising within these states are widely reported (Kugel et al., 2025) and are often described as unusually meaningful, emotionally intense, and sometimes ineffable (i.e. difficult to articulate) (Kallio-Mannila et al., 2025; Perkins et al.,2023; Timmermann et al., 2022; Vicentini et al., 2023). While these experiences share core characteristics with the “Aha!” moments described above, they often occur in a broader experiential context and are associated with more personally salient content and stronger subjective impact. Their content tends to centre on broad, self-relevant themes including identity, relationships, trauma, values, and existential meaning, and may be mystical or metaphysical in nature, involving abstract or emotionally complex material (Belser et al., 2017; Chopra & Letheby, 2024; Letheby, 2021; Watts et al., 2017). Importantly, psychedelic insights also appear likely to translate into psychological and behavioural change (Perkins et al., 2023). Supporting this, a recent systematic review reported that the degree of insight experienced during psychedelic sessions across clinical, experimental and naturalistic settings predicts subsequent therapeutic improvement, sometimes even more strongly than the traditional measure of “mystical experience” (Kugel et al., 2025). Together, these findings suggest that the subjective impact of insight—how compelling, meaningful, and intense it feels—may be central for understanding when it is experienced as belief-updating.

This raises two central questions for insight research: (1) To what extent do insight experiences differ across contexts (e.g. laboratory, everyday life, and psychedelic settings) in their phenomenological characteristics? (2) When insight-related belief change differs across contexts, is this difference driven by context per se, or by specific phenomenological dimensions (e.g., meaning, intensity, ineffability) that certain contexts tend to amplify?

From a theoretical perspective, recent work suggests that the phenomenology of insight may play a functional role in cognition. Specifically, the Eureka Heuristic account proposes that the feeling of insight acts as an adaptive metacognitive signal that helps evaluate and select among competing ideas (Laukkonen et al., 2022, 2023). According to this view, when a candidate idea is experienced as especially salient and coherent, it reaches awareness with a characteristic phenomenology—such as surprise, confidence, positive affect, and motivational drive—which may serve as a heuristic cue that the idea is important, trustworthy and worth endorsing.

Building on this perspective, it is useful to distinguish between core heuristic features of insight and appraised, context-sensitive qualities. Core features—such as surprise, confidence, positive affect, and perhaps motivational drive—are proposed to constitute the “Aha!” signal itself and may contribute to rapid commitment to an emerging interpretation. In contrast, qualities such as meaning, ineffability, and perceived transformative impact may depend more strongly on the content of the insight, the individual’s goals, and the context in which the insight occurs. This distinction is consistent with Tulver et al.’s (2025) proposal that mental breakthroughs vary along a continuum of phenomenological intensity and impact, depending partly on how central the restructured representation is within a person’s broader conceptual landscape. From this perspective, brief laboratory insights may involve relatively local restructuring, whereas everyday and psychedelic insights may more often involve self-relevant or higher-level representations. This provides a basis for expecting larger context differences in dimensions such as meaning, ineffability, and perceived belief change than in core “Aha!” features such as surprise, confidence, and pleasure.

Psychedelic contexts may be particularly likely to amplify these appraised qualities, given that the insights they often involve concern higher-level, self-relevant beliefs. One framework that may help explain this pattern is the REBUS model (RElaxed Beliefs Under Psychedelics; Carhart-Harris & Friston, 2019). Within this account, psychedelics are proposed to relax high-level priors (i.e. strongly held beliefs), reducing top-down constraints on perception and cognition and allowing novel interpretations to emerge more easily. These interpretations may be experienced as especially salient and intense, as emotional processing is less tightly constrained (Laukkonen et al., 2020; McGovern et al., 2024).

It would therefore be expected that phenomenological ratings for psychedelic insights—particularly intensity, meaning, ineffability and belief change—would be higher than those reported in everyday-life and laboratory contexts. More generally, reported belief change may depend less on context itself than on the phenomenological qualities that certain contexts tend to amplify, particularly meaning and intensity (Laukkonen et al., 2020, 2021, 2022).

The present study examined insight experiences across naturalistic contexts (psychedelic and everyday-life insights) and laboratory contexts (CRA and ambiguous-image tasks) using a within-subject, cross-sectional design. Participants rated insights across several phenomenological dimensions and reported the perceived impact of each insight on their beliefs. We examined context-related differences in insight phenomenology and tested whether specific phenomenological dimensions predicted perceived belief change beyond context alone.

Our aims were to (1) characterise context-related differences in insight phenomenology across naturalistic and laboratory contexts; (2) identify which phenomenological dimensions independently predict perceived belief change after accounting for context and participant-level variability. It was hypothesized that: (i) psychedelic insights would be rated higher in meaning, intensity, and ineffability, and show greater insight-related belief change, than everyday-life and laboratory insights; (ii) contextual differences between naturalistic and laboratory insight experiences were expected to vary across phenomenological dimensions, with larger differences anticipated for dimensions related to personal meaning and transformation (i.e., meaning, belief change) than for core “Aha!” features (i.e., surprise, confidence, pleasure, and drive); and (iii) within the naturalistic contexts, meaning would be the strongest positive predictor of perceived belief change, with intensity as a secondary predictor.

## 2 METHOD

Figure 1 provides a graphical overview of the study design, survey procedure, measures, and analytic approach.

**Figure 1.**
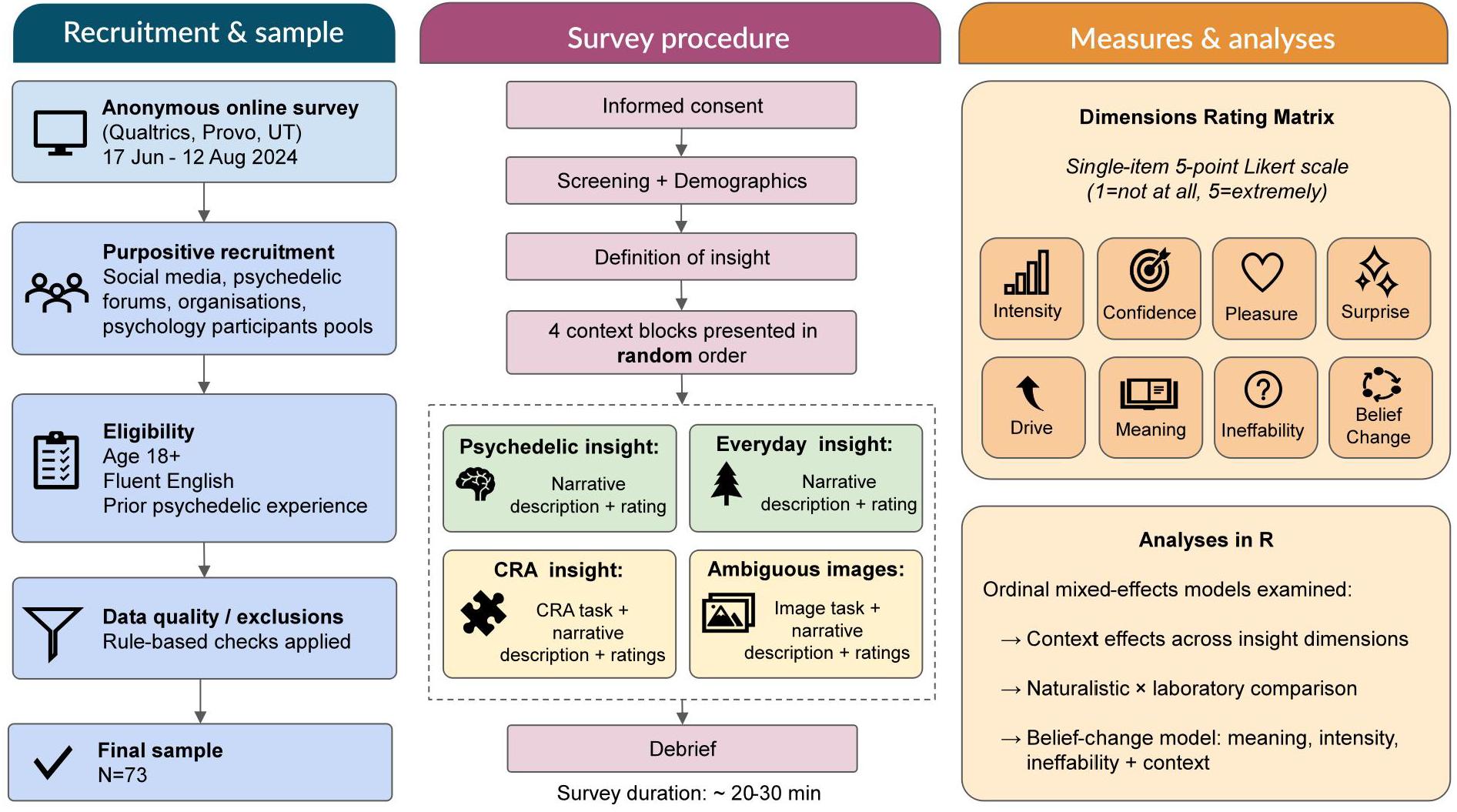
Overview of study design and procedure. **Note**. CRA = Compound Remote Associates. Green shading indicates naturalistic insight contexts; yellow shading indicates laboratory insight contexts.

### 2.1. Design and participants

This study employed a cross-sectional, within-subject design. Data were collected via an online survey in which participants reported insight experiences across four contexts (psychedelic, everyday life, CRA, and ambiguous image-based problem-solving). For each context, participants provided narrative descriptions and rated multiple phenomenological dimensions of insight. The study was managed on the Open Science Framework, with publicly available materials and analysis resources described in the Data Availability statement.

### 2.2. Procedure

After providing informed consent, participants completed a brief screening section assessing eligibility criteria, followed by demographic questions.

Participants were then provided with a definition of an insight (‘Aha!’) experience before completing four context blocks, presented in randomised order across participants.

In the two naturalistic contexts, participants recalled a personally meaningful psychedelic insight and a non-psychedelic everyday-life insight, provided narrative descriptions, and completed a set of phenomenological rating items. In the two laboratory contexts, participants completed CRA problems and ambiguous image tasks, then indicated whether they had experienced an “Aha!” moment. When an insight was reported, participants provided a brief description and completed the same rating items.

The survey concluded with a debriefing page and took approximately 20–30 minutes to complete.

### 2.3. Measures

#### Insight dimensions

To our knowledge, no single validated instrument currently captures the full set of experiential dimensions targeted here—particularly meaning, ineffability, and insight-related belief change—in a form that can be applied uniformly across both laboratory and naturalistic insight contexts. We therefore constructed a Dimensions Rating Matrix comprising single-item 5-point Likert-type ratings (1 = not at all; 5 = extremely), presented identically across contexts to enable within-subject comparisons.

The selected dimensions were derived from previous work on the phenomenology of insight and “Aha!” experiences, which has emphasised features such as suddenness/surprise, confidence or certainty, positive affect, and motivational drive (Danek et al., 2014; Danek & Wiley, 2017), as well as from naturalistic and psychedelic insight literature highlighting meaning, intensity, ineffability, and perceived belief change (Hill & Kemp, 2018; Kallio-Mannila et al., 2025; Perkins et al., 2023; Timmermann et al., 2021; Vicentini et al., 2023). Item wording was carefully considered to minimise ambiguity and maximise readability, and was piloted with a cohort of psychology students prior to the main study. For the problem-solving contexts, ratings were collected only when participants indicated that they had experienced an insight during the task.

Although belief change was measured using the same response format, it was treated as a key outcome in subsequent models testing phenomenological predictors of belief updating.

#### Narrative descriptions

For each context in which an insight was reported, participants provided an open-ended description of the insight experience in their own words. These descriptions were used to verify that responses were consistent with task instructions and eligibility criteria, but were not analysed as qualitative data in the present study.

### 2.4. Data preparation and exclusion criteria

Data preprocessing was conducted prior to all analyses using a rule-based pipeline implemented in R. A total of 257 individuals initiated the survey. Responses originating from survey preview or test links were removed. Participants were required to meet screening criteria (age ≥ 18 years, English proficiency, and prior psychedelic experience) and to provide complete insight ratings for both the psychedelic and everyday contexts. For the problem-solving tasks, completion rules were conditional: participants who reported not experiencing an insight were retained, whereas participants who reported an insight were required to complete all associated ratings for that task. Participants with incomplete ratings in any required section were excluded.

Additional exclusions were applied to ensure data quality and construct validity. Narrative responses were screened to verify compliance with task instructions and eligibility criteria and to identify responses containing safety-related content. Participants showing invalid response patterns (e.g., uniform responding across items) were also excluded. Given the study’s focus on classic psychedelic experiences, participants reporting cannabis-only psychedelic experiences were excluded. Following all preprocessing and exclusion steps, the final quantitative sample consisted of 73 participants.

### 2.5. Analyses

All quantitative analyses were conducted in R version 4.5.1 (R Core Team, 2025). Cumulative link mixed-effects models were fitted using the ordinal package (Christensen, 2023), and figures were produced using ggplot2 (Wickham, 2016). Insight ratings were measured on 5-point Likert-type scales and analysed as ordinal outcomes. Accordingly, cumulative link mixed-effects models (CLMMs) with a logit link function were used to model the ordered categorical data while accounting for the within-subject repeated-measures structure.

To examine context-related differences in insight phenomenology, separate CLMMs were fitted for each insight dimension (intensity, confidence, pleasure, surprise, meaning, ineffability, drive, and belief change). In each model, context (psychedelic, everyday life, CRA problem-solving, and image-based problem-solving) was included as a fixed effect, and participant was included as a random intercept to account for within-subject dependence. Everyday insights served as the reference category.

To examine whether the difference between naturalistic and laboratory insight experiences varied across phenomenological dimensions, we conducted an additional CLMM. For this analysis, psychedelic and everyday insights were combined into a naturalistic category, and CRA and image-based insights were combined into a laboratory category. The model included context type, dimension, and their interaction as fixed effects, with participant included as a random intercept. The interaction term was used to test whether the naturalistic–laboratory difference was larger for some dimensions than for others.

To examine phenomenological predictors of perceived belief change, an additional CLMM was fitted with belief change as the ordinal outcome, perceived meaning, intensity, and ineffability as fixed effects, and a random intercept for participant. Context (psychedelic vs. everyday) was included as a fixed effect to account for baseline differences between contexts, with everyday insights as the reference category. The primary belief-change model focused on psychedelic and everyday insights, as belief-change ratings in problem-solving contexts exhibited pronounced floor effects, limiting their suitability for primary inference. As a robustness check, the same model was refitted including all contexts (psychedelic, everyday, CRA, and image-based problem-solving), with results reported as a sensitivity analysis.

Model estimates are reported as log-odds (β) with corresponding standard errors and 95% confidence intervals. Statistical significance was evaluated using Wald tests for fixed effects. All tests were two-tailed with α = .05.

### 2.6. Ethics

The study was approved by the Southern Cross University Human Research Ethics Committee (Approval No. 2024/066). All participants provided informed consent prior to participation by initiating the online survey. Participation was anonymous, and no identifying information was collected. Given the sensitive nature of questions about psychedelic use, particular care was taken to ensure participant confidentiality. All data were de-identified and stored securely in accordance with university ethics requirements.

## 3. RESULTS

### 3.1. Demographic data

The final analytic sample comprised 73 participants. Most participants reported male gender (58.9%) and were aged 31–45 years (47.9%). Participants most commonly reported psychedelic insights occurring on psilocybin (32.9%), LSD (30.1%), or ayahuasca (23.3%), most often in personal/recreational settings (65.8%). Everyday insights were most frequently reported at home (41.1%) or in nature (21.9%). As expected given the survey design, ratings for the problem-solving contexts were available only when participants reported experiencing an insight, resulting in smaller sample sizes for the CRA (n = 46) and image-based (n = 53) contexts. Table 1 provides the frequencies and percentages for participant demographic and study characteristics.

**Table 1.** Sample characteristics (N = 73)

| Demographics | n | % | Demographics | n | % |
| --- | --- | --- | --- | --- | --- |
| *Age* |  |  | *Psychedelic substance* |  |  |
| 18–30 | 12 | 16.4 | Psilocybin | 24 | 32.9 |
| 31–45 | 35 | 47.9 | LSD | 22 | 30.1 |
| 46–60 | 17 | 23.3 | Ayahuasca | 17 | 23.3 |
| 61+ | 9 | 12.3 | Mescaline | 4 | 5.5 |
|  |  |  | DMT | 3 | 4.1 |
| *Gender* |  |  | Other | 3 | 4.1 |
| Male | 43 | 58.9 |  |  |  |
| Female | 29 | 39.7 | *Everyday setting* |  |  |
| Prefer not to say | 1 | 1.4 | Home | 30 | 41.1 |
| *Psychedelic setting* |  |  | Nature | 16 | 21.9 |
| Personal / recreational | 48 | 65.8 | While Traveling | 6 | 8.2 |
| Traditional / indigenous | 22 | 30.1 | Workplace | 5 | 6.8 |
| Clinical trial / therapy | 2 | 2.7 | Meditation | 4 | 5.5 |
| Prefer not to say | 1 | 1.4 | Other | 12 | 16.4 |

### 3.2. Descriptive statistics of insight ratings by context

Descriptive statistics for insight ratings across contexts and dimensions are presented in Figure 2, which summarises ordinal responses using median ratings. Across dimensions, psychedelic insights tended to receive higher ratings than everyday insights, while task-based insights (CRA and image-based) were generally rated lower. Everyday-life insights typically fell between these extremes. Full descriptive statistics and rating distributions by context and dimension are provided in Table S1.

**Figure 2.**
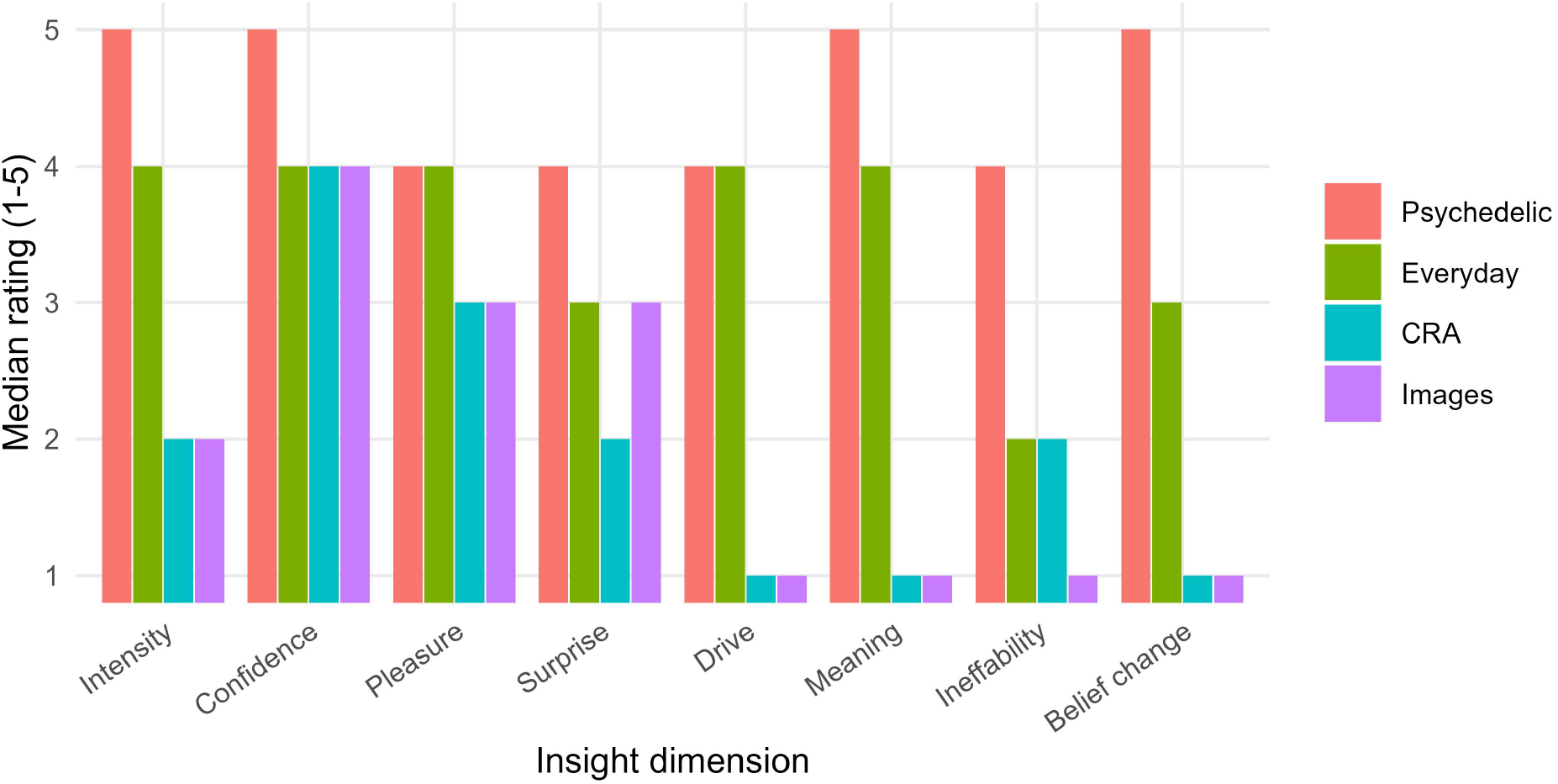
Median insight ratings across contexts and dimensions. **Note**. Bars indicate median ratings on a 1–5 ordinal scale.

### 3.3. Context effects on insight phenomenology

To examine whether the phenomenology of insight differed across contexts, separate cumulative link mixed-effects models were fitted for each insight dimension, with Context as a fixed effect and a random intercept for Participant. Ratings were treated as ordinal outcomes, and Everyday insights served as the reference category. Model estimates therefore reflect context-dependent shifts in the odds of reporting higher response categories. Figure 3 summarises context effects across dimensions (log-odds relative to everyday insights), and Supplementary Tables S2 and S2b report the corresponding estimates, confidence intervals, and test statistics.

**Figure 3.**
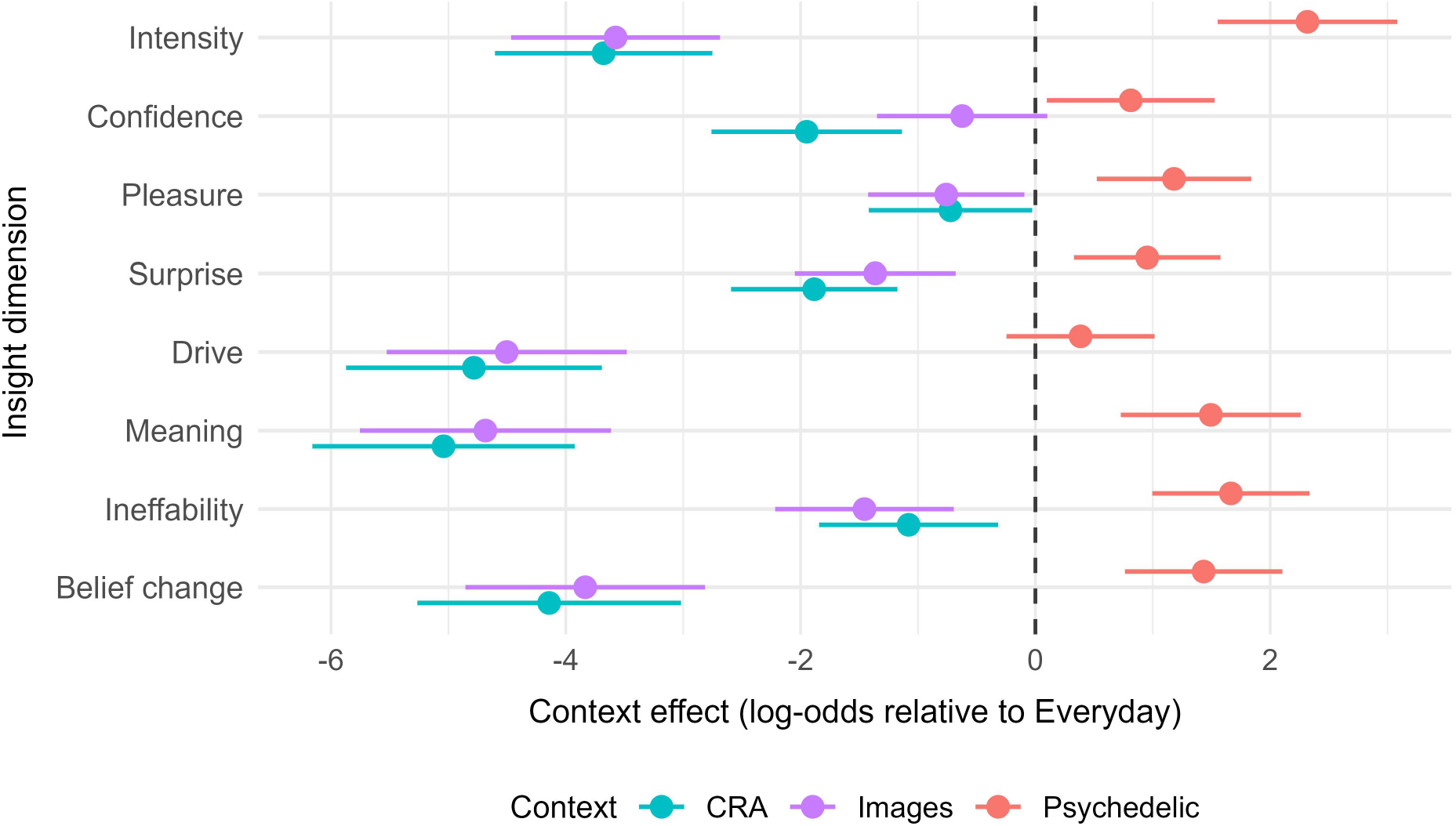
Context effects on insight dimensions relative to everyday insights. **Note**. Points indicate model estimates (log-odds) with 95% confidence intervals; the dashed line denotes the everyday reference (0).

Relative to everyday insights, psychedelic insights showed higher odds of stronger ratings across most dimensions. The largest effects were observed for intensity (β = 2.32, CI [1.55, 3.08], p < .001), ineffability (β = 1.67, CI [1.00, 2.34], p < .001), meaning (β = 1.49, CI [0.73, 2.26], p < .001), and belief change (β = 1.43, CI [0.76, 2.10], p < .001). Smaller positive effects were observed for pleasure, surprise, and confidence, while drive did not reliably differ between psychedelic and everyday insights (β = 0.38, 95% CI [−0.25, 1.02], p = .232).

In contrast, CRA and image-based insights were generally associated with lower ratings than everyday insights. The strongest reductions were observed for meaning, belief change, drive and intensity. For example, CRA insights showed substantially lower ratings for meaning (β = −5.04, 95% CI [−6.16, −3.92], p < .001), belief change (β = −4.14, 95% CI [−5.26, −3.02], p < .001), drive (β = −4.78, 95% CI [−5.87, −3.69], p < .001), and intensity (β = −3.68, 95% CI [−4.60, −2.75], p < .001). The image-based task showed a highly similar pattern of reductions across these dimensions (Table S2, S2b). Confidence was lower for CRA insights (β = −1.95, 95% CI [−2.76, −1.14], p < .001), but did not reliably differ between image-based and everyday insights (β = −0.62, 95% CI [−1.35, 0.10], p = .092).

### 3.4. Naturalistic vs laboratory differences across dimensions

To examine whether differences between naturalistic and laboratory insights varied across phenomenological dimensions, a cumulative link mixed-effects model including a ContextType × Dimension interaction was compared to a model without the interaction. Full model comparison results and fixed-effect estimates are provided in Tables S3a and S3b, respectively. The interaction significantly improved model fit (χ^2^(7) = 304.92, p < .001), indicating that the magnitude of the naturalistic–laboratory difference depended on the dimension being rated.

All dimensions were rated higher in naturalistic than laboratory contexts, but the magnitude of this difference varied substantially across dimensions (Table 2). The largest effects were observed for meaning (β = −5.38, 95% CI [−5.95, −4.81]) and belief change (β = −5.04, 95% CI [−5.68, −4.40]), followed by drive (β = −4.56, 95% CI [−5.12, −3.99]) and intensity (β = −3.38, 95% CI [−3.87, −2.90]). Moderate differences were observed for ineffability and surprise, whereas confidence and pleasure showed comparatively smaller differences. Importantly, this analysis compares naturalistic and laboratory contexts at a broad level and does not isolate psychedelic-specific effects, as psychedelic and everyday insights are combined within the naturalistic category.

**Table 2.** Naturalistic vs laboratory differences in insight dimensions.

| Dimension | $\beta$ (log-odds) | 95% CI | p |
| --- | --- | --- | --- |
| Meaning | -5.38 | [-5.95, -4.81] | <.001 |
| Belief Change | -5.04 | [-5.68, -4.40] | <.001 |
| Drive | -4.56 | [-5.12, -3.99] | <.001 |
| Intensity | -3.38 | [-3.87, -2.90] | <.001 |
| Ineffability | -2.11 | [-2.62, -1.60] | <.001 |
| Surprise | -1.94 | [-2.40, -1.47] | <.001 |
| Confidence | -1.17 | [-1.66, -0.68] | <.001 |
| Pleasure | -1.14 | [-1.60, -0.69] | <.001 |

**Note.** Values are log-odds ( $\beta$ ) from a cumulative link mixed-effects model. Negative values indicate higher ratings in naturalistic compared to laboratory contexts. Dimensions are ordered by effect size.

### 3.5. Predictors of perceived belief change

To examine whether phenomenological features of insight predicted perceived belief change within the two naturalistic contexts, a cumulative link mixed-effects model was fitted with belief change as the ordinal outcome and meaning, intensity, ineffability and context as fixed effects. Participant was included as a random intercept, and everyday insights served as the reference category. The primary model focused on psychedelic and everyday insights because belief-change ratings in problem-solving contexts showed pronounced floor effects. An all-context model was conducted as a sensitivity analysis. Within psychedelic and everyday contexts, perceived meaning was the strongest predictor of belief change (β = 1.57, CI [0.90, 2.25], p < .001). Intensity was also positively associated with belief change, although with a smaller effect size (β = 0.66, CI [0.12, 1.20], p = .017). Ineffability showed an additional independent association with belief change (β = 0.41, CI [0.05, 0.77], p = .024). After accounting for these phenomenological predictors, context (psychedelic vs. everyday) was not reliably associated with belief change (β = 0.34, CI [−0.46, 1.15], p = .406), suggesting that differences in belief change between psychedelic and everyday insights were largely accounted for by differences in perceived meaning, intensity, and ineffability rather than by context per se. Full model estimates are reported in Table 3.

**Table 3.** Predictors of Perceived Belief Change (Everyday and Psychedelic Contexts)

| Term | $\beta$ | SE | z | p | 95% CI |
| --- | --- | --- | --- | --- | --- |
| Context (Psychedelic vs. Everyday) | 0.34 | 0.41 | 0.83 | 0.41 | [-0.46, 1.15] |
| Meaning | 1.57 | 0.34 | 4.58 | < .001 | [0.90, 2.25] |
| Ineffability | 0.41 | 0.18 | 2.26 | 0.02 | [0.05, 0.77] |
| Intensity | 0.66 | 0.28 | 2.38 | 0.02 | [0.12, 1.20] |

**Note.** Values are log-odds ( $\beta$ ) from a cumulative link mixed-effects model with a logit link. 95% CIs are Wald intervals. Everyday is the reference for Context. Random intercept: Participant.

A sensitivity analysis including all contexts showed a highly similar pattern of results. Meaning remained the strongest predictor of belief change (β = 1.26, CI [0.82, 1.71], p < .001), with additional positive associations for intensity (β = 0.70, CI [0.27, 1.14], p = .002) and ineffability (β = 0.37, CI [0.08, 0.66], p = .013). None of the context contrasts relative to everyday insights were reliably associated with belief change after adjusting for phenomenological predictors (Table S4).

Overall, belief change was most strongly associated with perceived meaning of the insight, with smaller but reliable contributions from intensity and ineffability. These associations were observed within psychedelic and everyday contexts and generalised across contexts in sensitivity analyses, indicating that perceived belief change was more closely related to phenomenological qualities of insight than to the context in which the insight occurred.

## 4. DISCUSSION

The present study examined the phenomenology of insight across psychedelic, everyday-life, and laboratory contexts. The aim was to clarify (i) how insight experiences differ across contexts and (ii) which phenomenological dimensions are associated with perceived belief change. We found that insight ratings varied systematically across contexts but that these differences were not uniform across dimensions. In particular, dimensions related to personal significance (e.g. meaning and belief change) showed the largest differences between naturalistic and laboratory contexts. In contrast, dimensions typically associated with the core “Aha!” experience such as confidence and pleasure, showed comparatively smaller differences. Additionally, perceived belief change was most strongly associated with the perceived meaningfulness of the insight, rather than with context alone.

Consistent with our first hypothesis, psychedelic insights were rated higher than everyday insights on several dimensions, particularly intensity, meaning, ineffability, and belief change. These findings align with the idea that psychedelic experiences are characterised by increased subjective intensity, salience, and perceived significance of one’s own thoughts and feelings (McGovern et al., 2024; Laukkonen et al., 2020). Everyday insights typically showed intermediate ratings, whereas insights arising in laboratory tasks (CRA and ambiguous images) were associated with lower ratings across most dimensions.

These results support the view that laboratory paradigms capture a narrower range of insight experience. While such tasks reliably elicit “Aha!” moments, they appear to only partially capture aspects of insight related to personal relevance, meaning, and perceived impact on beliefs. Importantly, however, laboratory insights still exhibited the characteristic phenomenological profile associated with the “Aha!” experience, indicating that they successfully capture core features of insight.

A central aim of the study was to examine whether some dimensions of insight are relatively stable across contexts, while others are more context-sensitive. In line with our second hypothesis, the difference in rating between naturalistic (psychedelic and everyday) and laboratory (CRA and ambiguous images) contexts varied across dimensions. Meaning and belief change showed the largest differences, followed by drive and intensity, whereas confidence and pleasure showed comparatively smaller differences.

This pattern is consistent with the distinction proposed earlier between core heuristic features of insight and more appraised, context-dependent qualities. Dimensions such as confidence, pleasure, and surprise—often considered central components of the “Aha!” experience—showed relatively smaller context effects, suggesting that they may reflect a more general metacognitive signal of insight. Within the Eureka Heuristic framework (Laukkonen et al., 2023), these features can be understood as cues that an emerging idea has reached sufficient salience and coherence to be accepted and endorsed. In contrast, meaning and belief change showed significantly larger differences between contexts, supporting the view that these dimensions depend more strongly on the content and personal relevance of the insight. This is consistent with the fact that insights in everyday and psychedelic contexts often concern self-relevant or emotionally significant material, whereas laboratory tasks typically involve problems with limited personal importance.

Interestingly, drive – conceptualised as “motivation to act” and hypothesised to be a core insight feature – showed large differences between naturalistic and laboratory contexts, comparable to those of meaning and belief change. Motivational drive was originally identified as a relevant insight dimension in qualitative analyses of participants’ descriptions of insight experiences in laboratory paradigms (Danek et al., 2014). However, when examined quantitatively within similar paradigms, drive did not reliably predict “Aha!” ratings once individual differences were accounted for, suggesting that it may not constitute a core feature of the Aha experience (Danek & Wiley, 2017). The present findings extend previous work by showing that drive differs substantially across contexts. The fact that drive was rated higher in naturalistic than laboratory contexts suggests that its significance may depend on the perceived relevance of the insight. While laboratory insights may elicit a feeling of solution or correctness, they may be less likely to generate a broader motivational response. In contrast, insights in everyday and psychedelic contexts, which often involve personally meaningful or self-relevant content, may be more likely to produce an urge to act on the novel idea.

Consistent with our third hypothesis, meaning emerged as the strongest predictor of belief change within naturalistic contexts, with additional contributions from intensity and ineffability. Importantly, once these phenomenological dimensions were included in the model, context (psychedelic vs. everyday) was not reliably associated with belief change. This indicates that differences in perceived belief updating across contexts are better explained by differences in how insights are experienced than by context alone. This finding supports the idea that meaning might play a central role in determining whether an insight is integrated into existing belief structures. Regardless of context, insights that are experienced as meaningful may be more likely to be treated as relevant, increasing their likelihood of influencing beliefs. Intensity may further contribute by increasing the salience or perceived strength of an insight, while ineffability may characterise experiences that are difficult to integrate into existing ways of understanding. On this interpretation, ineffability may not itself drive belief change but may instead mark experiences that are more likely to be perceived as profound and relevant. Importantly, the present data cannot determine whether participants’ insights were accurate, or whether perceived belief change translated into longer-term changes in beliefs or behaviour. The findings therefore should be interpreted as evidence of perceived belief relevance rather than objective belief revision.

The present findings highlight both the strengths and limitations of laboratory-based approaches to studying insight. Laboratory paradigms successfully capture key features of the “Aha!” experience, including surprise, confidence, and positive affect. However, they appear to underrepresent dimensions related to personal meaning, belief updating, and motivational impact. As a result, conclusions about the functional role of insight based solely on laboratory paradigms may underestimate its potential impact on belief systems and behaviour. This underscores the importance of examining context-sensitive, appraised dimensions in naturalistic settings.

## 5. LIMITATIONS AND FUTURE DIRECTIONS

Several limitations of this study should be considered. First, the study relies on retrospective self-report, which may be influenced by memory biases and post-hoc reinterpretation. In particular, ratings of meaning and belief change may reflect participants’ current interpretations of the experience rather than its original phenomenological properties. Second, each phenomenological dimension was assessed using a single-item measure. While this approach allows for a consistent assessment across multiple contexts, single-item measures may be more susceptible to measurement error and may not fully capture the complexity of each construct. Third, the cross-sectional design limits causal inference. Although meaning, intensity, and ineffability were associated with perceived belief change, the direction of these relationships cannot be determined. Fourth, because eligibility required participants to have experienced an insight while under the effects of a psychedelic substance, the sample does not represent psychedelic users who do not report such insights, or individuals who experience insights in everyday or laboratory contexts but have not had psychedelic insights. Finally, although gender was recorded descriptively, the study was not designed to examine gender differences in insight phenomenology.

Future work should therefore examine insight across contexts using longitudinal and multi-method designs. Longitudinal studies could test whether dimensions such as meaning, intensity, and ineffability predict longer-term changes in beliefs, behaviour, and wellbeing. Future research should also evaluate and validate the dimensional structure used here with multi-item scales and factor-analytic approaches. Finally, qualitative analyses of insight narratives could clarify how different forms of insight content, such as self-related, relational, or existential insights, relate to specific phenomenological dimensions, and whether additional dimensions emerge in naturalistic and psychedelic contexts.

## 6. CONCLUSION

In summary, the present study shows that insight experiences share a common phenomenological pattern across contexts, but vary substantially in their perceived meaning, belief change, and motivational impact. The findings suggest that while laboratory paradigms capture key features of the “Aha!” experience, they may underrepresent aspects of insight that are most closely linked to belief updating. The findings also suggest that the impact of insight depends less on the context in which it occurs, and more on how it is experienced, particularly in terms of its perceived meaning.

## CRediT authorship contribution statement

**Ambra Pogliani**: Conceptualization, Methodology, Formal analysis, Data curation, Visualization, Writing – original draft, Writing – review and editing. **Alexandra Zachary**: Conceptualization, Methodology, Investigation, Data curation, Resources, Writing – review and editing. **Lena Hall**: Investigation, Writing – review and editing. **Jason M. Tangen**: Investigation, Writing – review and editing. **Jonathan Mond**: Conceptualization, Supervision, Writing – review and editing. **Ruben E.Laukkonen**: Conceptualization, Methodology, Supervision, Writing – review and editing.

## Ethics approval

The study was approved by the Southern Cross University Human Research Ethics Committee (Approval No. 2024/066; approval date: 11/06/2024). All participants provided informed consent prior to participation.

## Declaration of competing interest

The authors declare that they have no known competing financial interests or personal relationships that could have appeared to influence the work reported in this paper.

## Funding

This research did not receive any specific grant from funding agencies in the public, commercial, or not-for-profit sectors.

## Acknowledgements

The authors thank the participants who contributed their time to this study. We also thank the psychology students who provided feedback during piloting of the insight dimension items.

## Data availability statement

Study materials, de-identified quantitative data, and R analysis code are available on the Open Science Framework at https://osf.io/9dvt8. Raw free-text responses are not publicly available due to ethical and privacy restrictions, as they may contain sensitive or potentially identifiable participant-generated content.

## Declaration of generative AI and AI-assisted technologies in the writing process

During the preparation of this work, the authors used ChatGPT to support language refinement, structural editing, clarity of expression, and troubleshooting of R code for statistical analysis and data visualisation. All AI-assisted text was manually reviewed and edited by the authors, and the code, model outputs, and interpretation of results were independently verified. The authors take full responsibility for the final content of the article.

## Notes

https://osf.io/9dvt8

### Competing Interest Statement

The authors have declared no competing interest.

### Summary of Updates

Figure 1's quality in PDF version of manuscript.

