## Supplementary material for "One-shotted: Psychedelic insights are uniquely intense, meaningful, and ineffable"

**Table S1**

*Descriptive statistics for insight ratings by context and dimension*

| **Dimension** | **Psychedelic** | **Everyday** | **CRA** | **Images** |
| --- | --- | --- | --- | --- |
| Intensity | 5 (1) | 4 (1) | 2 (2) | 2 (2) |
| Confidence | 5 (1) | 4 (1) | 4 (1) | 4 (1) |
| Pleasure | 4 (2) | 4 (2) | 3 (2) | 3 (2) |
| Surprise | 4 (2) | 3 (1) | 2 (1) | 3 (1) |
| Meaning | 5 (0) | 4 (1) | 1 (1) | 1 (1) |
| Ineffability | 4 (3) | 2 (2) | 2 (1) | 1 (1) |
| Drive | 4 (2) | 4 (2) | 1 (1) | 1 (1) |
| Belief change | 5 (1) | 3 (3) | 1 (0) | 1 (0) |

**Note**. Values are medians with interquartile ranges in parentheses. Ratings were provided on 5-point Likert-type scales (1 = not at all; 5 = extremely). Ns vary across contexts because ratings for problem-solving tasks were collected only when participants reported experiencing an insight (psychedelic and everyday, N = 73; CRA, n = 46; images, n = 53).

**Table S2**

*Context effects from dimension-specific ordinal mixed-effects models*

| Dimension | Psychedelic vs. Everyday | CRA vs. Everyday | Images vs. Everyday |
| --- | --- | --- | --- |
| Intensity | 2.32 [1.55, 3.08] | -3.68 [-4.60, -2.75] | -3.58 [-4.47, -2.69] |
| Confidence | 0.81 [0.10, 1.53] | -1.95 [-2.76, -1.14] | -0.62 [-1.35, 0.10] |
| Pleasure | 1.18 [0.52, 1.84] | -0.72 [-1.42, -0.03] | -0.76 [-1.42, -0.09] |
| Surprise | 0.95 [0.33, 1.58] | -1.88 [-2.59, -1.18] | -1.36 [-2.05, -0.68] |
| Drive | 0.38 [-0.25, 1.02] | -4.78 [-5.87, -3.69] | -4.50 [-5.53, -3.48] |
| Meaning | 1.49 [0.73, 2.26] | -5.04 [-6.16, -3.92] | -4.68 [-5.75, -3.62] |
| Ineffability | 1.67 [1.00, 2.34] | -1.08 [-1.84, -0.32] | -1.46 [-2.22, -0.70] |
| Belief Change | 1.43 [0.76, 2.10] | -4.14 [-5.26, -3.02] | -3.83 [-4.86, -2.81] |

**Note**. Values are log-odds (logit scale) relative to Everyday (reference). Cells show Estimate [95% Wald CI]. Positive values indicate higher odds of higher ratings compared to Everyday; negative values indicate lower odds.

**Table S2b**

*Full fixed-effect output (CLMM)*

| Dimension | Context | β | SE | z | 95% CI | p |
| --- | --- | --- | --- | --- | --- | --- |
| Intensity | Psychedelic | 2.32 | 0.39 | 5.94 | [1.55, 3.08] | < .001 |
| Intensity | CRA | -3.68 | 0.47 | -7.78 | [-4.60, -2.75] | < .001 |
| Intensity | Images | -3.58 | 0.45 | -7.87 | [-4.47, -2.69] | < .001 |
| Confidence | Psychedelic | 0.81 | 0.36 | 2.23 | [0.10, 1.53] | 0.026 |
| Confidence | CRA | -1.95 | 0.41 | -4.71 | [-2.76, -1.14] | < .001 |
| Confidence | Images | -0.62 | 0.37 | -1.69 | [-1.35, 0.10] | 0.092 |
| Pleasure | Psychedelic | 1.18 | 0.34 | 3.52 | [0.52, 1.84] | < .001 |
| Pleasure | CRA | -0.72 | 0.36 | -2.04 | [-1.42, -0.03] | 0.042 |
| Pleasure | Images | -0.76 | 0.34 | -2.24 | [-1.42, -0.09] | 0.025 |
| Surprise | Psychedelic | 0.95 | 0.32 | 2.99 | [0.33, 1.58] | 0.003 |
| Surprise | CRA | -1.88 | 0.36 | -5.21 | [-2.59, -1.18] | < .001 |
| Surprise | Images | -1.36 | 0.35 | -3.90 | [-2.05, -0.68] | < .001 |
| Drive | Psychedelic | 0.38 | 0.32 | 1.20 | [-0.25, 1.02] | 0.232 |
| Drive | CRA | -4.78 | 0.56 | -8.60 | [-5.87, -3.69] | < .001 |
| Drive | Images | -4.50 | 0.52 | -8.63 | [-5.53, -3.48] | < .001 |
| Meaning | Psychedelic | 1.49 | 0.39 | 3.82 | [0.73, 2.26] | < .001 |
| Meaning | CRA | -5.04 | 0.57 | -8.85 | [-6.16, -3.92] | < .001 |
| Meaning | Images | -4.68 | 0.55 | -8.59 | [-5.75, -3.62] | < .001 |
| Ineffability | Psychedelic | 1.67 | 0.34 | 4.87 | [1.00, 2.34] | < .001 |
| Ineffability | CRA | -1.08 | 0.39 | -2.78 | [-1.84, -0.32] | 0.006 |
| Ineffability | Images | -1.46 | 0.39 | -3.75 | [-2.22, -0.70] | < .001 |
| Belief Change | Psychedelic | 1.43 | 0.34 | 4.19 | [0.76, 2.10] | < .001 |
| Belief Change | CRA | -4.14 | 0.57 | -7.23 | [-5.26, -3.02] | < .001 |
| Belief Change | Images | -3.83 | 0.52 | -7.36 | [-4.86, -2.81] | < .001 |

**Note**. Estimate = log-odds; SE = standard error; z = Wald statistic; p = Wald p-value.

**Table S3a**

*Omnibus ContextType × Dimension interaction test*

| Model | Parameters | AIC | logLik | LR | df | p |
| --- | --- | --- | --- | --- | --- | --- |
| h2_main_model | 13 | 5138.18 | -2556.09 | NA |  |  |
| h2_interaction_model | 20 | 4847.26 | -2403.63 | 304.92 | 7 | < .001 |

**Note.** Nested cumulative link mixed-effects models were compared using a likelihood-ratio test. The key test is whether the interaction model fits better than the main-effects model.

### **Table S3b**

*Full fixed-effect output for the interaction model*

| Term | Estimate | SE | z | 95% CI | p |
| --- | --- | --- | --- | --- | --- |
| ContextType: Laboratory vs Naturalistic | -3.38 | 0.25 | -13.61 | [-3.87, -2.90] | < .001 |
| Dimension: Confidence vs Intensity | 0.75 | 0.23 | 3.28 | [0.30, 1.20] | < .001 |
| Dimension: Pleasure vs Intensity | -0.63 | 0.22 | -2.81 | [-1.06, -0.19] | < .001 |
| Dimension: Surprise vs Intensity | -0.83 | 0.22 | -3.77 | [-1.26, -0.40] | < .001 |
| Dimension: Drive vs Intensity | -0.24 | 0.22 | -1.10 | [-0.68, 0.19] | 0.27 |
| Dimension: Meaning vs Intensity | 0.96 | 0.24 | 4.06 | [0.50, 1.43] | < .001 |
| Dimension: Ineffability vs Intensity | -1.97 | 0.23 | -8.51 | [-2.43, -1.52] | < .001 |
| Dimension: Belief change vs Intensity | -0.44 | 0.23 | -1.94 | [-0.88, 0.00] | 0.05 |
| Interaction: Laboratory × Confidence | 2.21 | 0.35 | 6.40 | [1.54, 2.89] | < .001 |
| Interaction: Laboratory × Pleasure | 2.24 | 0.33 | 6.76 | [1.59, 2.89] | < .001 |
| Interaction: Laboratory × Surprise | 1.45 | 0.33 | 4.37 | [0.80, 2.09] | < .001 |
| Interaction: Laboratory × Drive | -1.17 | 0.36 | -3.24 | [-1.88, -0.46] | < .001 |
| Interaction: Laboratory × Meaning | -1.99 | 0.36 | -5.57 | [-2.70, -1.29] | < .001 |
| Interaction: Laboratory × Ineffability | 1.27 | 0.35 | 3.67 | [0.59, 1.95] | < .001 |
| Interaction: Laboratory × Belief change | -1.66 | 0.39 | -4.23 | [-2.43, -0.89] | < .001 |

**Note.** Estimate = log-odds; SE = standard error; z = Wald statistic; p = Wald p-value. Naturalistic and Intensity are the reference categories.

**Table S4**

*Predictors of belief change (all contexts).*

| Term | β | SE | z | p | 95% CI |
| --- | --- | --- | --- | --- | --- |
| Context: Psychedelic (vs. Everyday) | 0.24 | 0.38 | 0.64 | 0.53 | [-0.50, 0.99] |
| Context: CRA (vs. Everyday) | -0.95 | 0.70 | -1.37 | 0.17 | [-2.31, 0.41] |
| Context: Images (vs. Everyday) | -0.30 | 0.65 | -0.46 | 0.64 | [-1.57, 0.97] |
| Meaning | 1.26 | 0.23 | 5.56 | < .001 | [0.82, 1.71] |
| Ineffability | 0.37 | 0.15 | 2.49 | 0.01 | [0.08, 0.66] |
| Intensity | 0.70 | 0.22 | 3.15 | < .001 | [0.27, 1.14] |

**Note**. Sensitivity model includes Everyday, Psychedelic, CRA, and Images contexts. Everyday is the reference. Random intercept: Participant.
